# X-Linked Histone H3K27 Demethylase Kdm6a Regulates Sexually Dimorphic Differentiation of Hypothalamic Neurons

**DOI:** 10.1101/2021.06.03.446961

**Authors:** Lucas E. Cabrera Zapata, Carla D. Cisternas, Camila Sosa, Maria Angeles Arevalo, Luis Miguel Garcia-Segura, María Julia Cambiasso

## Abstract

Several X-linked genes are involved in neuronal differentiation and may contribute to the generation of sex dimorphisms in brain. Previous results showed that XX hypothalamic neurons grow faster, have longer axons, and exhibit higher expression of the neuritogenic gene *neurogenin 3* (*Ngn3*) than XY before perinatal masculinization. Here we evaluated the participation of candidate X-linked genes in the development of these sex differences, focusing mainly on *Kdm6a*, a gene encoding for an H3K27 demethylase with functions controlling gene expression genome-wide. We established hypothalamic neuronal cultures from wild-type or transgenic Four Core Genotypes mice, a model that allows evaluating the effect of sex chromosomes independently of gonadal type. X-linked genes *Kdm6a*, *Eif2s3x* and *Ddx3x* showed higher expression in XX compared to XY neurons, regardless of gonadal sex. Moreover, *Kdm6a* expression pattern with higher mRNA levels in XX than XY did not change with age at E14, P0, and P60 in hypothalamus or under 17β-estradiol treatment in culture. Kdm6a pharmacological blockade by GSK-J4 reduced the expression of neuritogenic genes *Neurod1*, *Neurod2* and *Cdk5r1* in both sexes equally, while a sex-specific effect was observed on *Ngn3* levels, with a decrease in XX and an increase in XY neurons. Finally, both Kdm6a inhibition and its downregulation using siRNA reduced axonal length only in female neurons, abolishing the sex differences observed in control conditions. Altogether, these results point to *Kdm6a* as a key mediator of the higher axogenesis and *Ngn3* expression observed in XX neurons before critical period of brain masculinization.

## INTRODUCTION

The brain is a sexually dimorphic organ, along with many other organs and tissues besides the gonads. These sex differences are found on a wide range of levels, both in the biochemical and architectural nature of certain brain regions and, consequently, in the physiological and behavioral responses controlled by such dimorphic regions, as well as in the susceptibility to certain neurodevelopmental, psychiatric, and neurodegenerative diseases, such as autism spectrum disorders, dyslexia, depression and anxiety disorders, attention deficit hyperactivity disorder, schizophrenia, and dementia, among others [1, 2]. Two major factors are currently known to contribute to the setting of sexual dimorphisms in the brain during development: (1) a sex-specific trophic environment due to differences in gonadal hormones secretion and (2) a distinct genetic and epigenetic pattern for males and females generated by differences in the expression of X and Y chromosomes-linked genes [3–5].

X and Y chromosomes have undergone a divergent evolutionary process, determining that they currently encode mostly dissimilar genetic information and are subject to different epigenetic regulations. As a result, these chromosomes are no longer capable of recombining during meiosis along most of their length except for the pseudoautosomal regions (PARs), the small regions of sequence identity at the termini of both X and Y. Therefore, the intermediary regions between the PARs that cannot recombine, called the non-PAR of the X and the male specific region of the Y (MSY), encompass most of both chromosomes and encode genes that are X- and Y-specific, respectively [6, 7]. Although X-linked genes are present in both sexes, the existence of two X in every cell of females and only one in males generates a “dosage difference” in the copy number of virtually all of these genes between the sexes. This imbalance is compensated during development by the X chromosome inactivation mechanism (XCI), which involves the transcriptional silencing of one of the two Xs in each cell of an XX embryo, thus defining an inactive (Xi) and an active (Xa) X chromosome that will be inherited through the successive mitotic divisions to all the cells that finally shape that individual [8–11]. However, XCI does not lead to complete repression of all genes in the Xi, but some “escape” inactivation and are therefore expressed from both Xa and Xi [12–15].

*Kdm6a* is an XCI “escapee” located on the non-PAR of the X that encodes for the lysine demethylase 6a (Kdm6a) or Utx enzyme, a member of the Kdm6 subfamily of histone 3 (H3) demethylases that remove di-(me2) and trimethyl (me3) groups on lysine (K) at position 27 (H3K27me2/me3) [16]. H3K27me2/me3 are epigenetic marks known to repress gene expression, so their removal by Kdm6 demethylases promotes transcription [17–19]. Along with Kdm6a, the other two members of the Kdm6 subfamily are lysine demethylase 6b (Kdm6b) or Jmjd3, encoded on autosome 11, and lysine demethylase 6c (Kdm6c) or Uty, a Kdm6a paralogue encoded on the MSY [16, 19, 20]. Kdm6a plays a key role in cell fate determination and cellular identity during development by controlling pluripotency and lineage-specific sets of genes [21, 22]. During brain development, Kdm6a is involved in neurogenesis promotion, determining the neural stem cell status and modulating the subsequent differentiation of these pluripotent cells into neurons and glia [23–26]. Regarding neuronal differentiation, it has been shown that Kdm6a deletion prevents the proper development and maturation of neurons, leading to a repression of genes required for neuritogenesis and synaptogenesis (such as *CREB5*, *CAMK2A*, *CKB*, *ASIC1*, and *ASCL1*, among many others), defects and immaturity in dendritic arborization and synapse formation, failures in electrophysiological activity, increased expression of anxious behaviors, and deficits in spatial learning and memory [27, 28]. Finally, *Kdm6a* constitutive mutations with loss of function lead to Kabuki syndrome, a congenital disorder characterized by intellectual disability accompanied by growth retardation and skeletal, cardiac, and facial abnormalities, among other manifestations [29–31].

The use of the Four Core Genotypes (FCG) transgenic mice allows independent evaluation of sex hormones and X and Y chromosomes effects on sexual differentiation. FCG model conjugates the deletion of the testis-determining *Sry* gene from the Y chromosome [32] with the reinsertion of this gene into autosome 3 [33]. This way, it was possible to unlink the inheritance of the Y chromosome from the inheritance of *Sry* and testis differentiation, obtaining four distinct genotypes: XX males (XXm) and females (XXf) and XY males (XYm) and females (XYf). Our previous results using FCG mice showed that sex chromosome complement (XY/XX) determines a sexually dimorphic expression of the proneural basic Helix-Loop-Helix transcription factor *neurogenin 3* (*Ngn3*), with XX hypothalamic neurons presenting higher levels of *Ngn3* mRNA than XY neurons, independently of gonadal sex [34]. In turn, XX neurons showed faster growth and maturation *in vitro* than XY neurons, a characteristic that was dependent on the higher *Ngn3* expression in XX cultures [34, 35]. These and other results [36–41] point to the importance of sex chromosome complement in the sexual differentiation of the brain, regulating the expression of autosomal genes involved in neurodevelopment and modulating neuronal differentiation in a sex-specific way independently of sex hormones. However, although it is clear that sex chromosomes determined the sexual dimorphisms observed in *Ngn3* expression and neuronal growth and differentiation, the identity of particular X and Y-linked genes taking part in these processes remains unknown. Therefore, in the present study we aimed to evaluate specific X-linked genes participation in the sexually dimorphic differentiation of hypothalamic neurons, focusing on *Kdm6a* as a genome-wide regulator of gene expression that escapes XCI and has a significant role in neurogenesis and neuritogenesis.

## MATERIALS AND METHODS

### Animals

FCG transgenic mice generated from MF1 strain (MF1-FCG mice, kindly donated by Dr. Paul Burgoyne, National Institute for Medical Research, London, UK), MF1 wild-type mice (Harlan Laboratories Inc., USA), and CD1 wild-type mice were used. MF1-FCG and MF1 wild-type mice were bred at the Instituto M. y M. Ferreyra (INIMEC-CONICET-Universidad Nacional de Córdoba, Córdoba, Argentina), whereas CD1 mice were obtained from colonies bred at both the Instituto M. y M. Ferreyra and the Instituto Cajal (CSIC, Madrid, Spain). Embryos used in FCG cultures were produced by breeding transgenic MF1-FCG XYm mice with wild-type MF1 females. Animals were kept in specific pathogen free (SPF) conditions in individually ventilated cages until crosses were made to obtain pregnant females for experiments, at which time animals were placed in open cages. In all cases, the animals received water and food *ad libitum* and were kept in controlled macroenvironmental conditions of temperature at 23 °C and a 12 h light/12 h dark periodic cycle. Procedures for care, welfare and proper use of all experimental animals were approved and controlled by the Institutional Animal Care and Use Committee (CICUAL) of the Instituto M. y M. Ferreyra, following national and international regulations, and were in accordance with the Consejería del Medio Ambiente y Territorio (Comunidad de Madrid, Ref. PROEX 200/14), the European Commission (86/609/CEE and 2010/63/UE), and the Spanish Government Directive (R.D.1201/2005) guidelines. FCG animals genotyping was performed by Polymerase Chain Reaction (PCR) as previously described in Cisternas *et al*. [41].

### Hypothalamic neuronal cultures and cell treatments

FCG or CD1 wild-type embryos at 14 days of gestation (E14, defining E0 as the day of the vaginal plug) were used to establish primary hypothalamic neuronal cultures. Donor embryos age was specifically selected with the purpose of avoiding exposure of neurons to the peak in gonadal testosterone secretion during *in utero* development, which occurs in male mice around E17 [42]. Pregnant mice were sacrificed by cervical dislocation under CO_2_ anesthesia, and embryos were dissected from the uterus. Neurons were cultured separately according to sex (by observing the presence/absence of the spermatic artery in developing gonads) or genotype (by PCR) of embryos. The ventromedial hypothalamic region was dissected out and stripped off the meninges. Blocks of tissue were incubated for 15 min at 37 °C with 0.5% trypsin (Gibco, USA) and then washed three times with Ca^2+^/Mg^2+^-free Hank’s Buffered Salt Solution. Finally, tissue was mechanically dissociated to single cells in 37 °C warm culture medium and cells were seeded. The medium was phenol red-free Neurobasal (Gibco) to avoid “estrogen-like effects” [43] and was supplemented with B-27, 0.043% L-alanyl-L-glutamine (GlutaMAX-I), 100 U/ml penicillin, and 100 mg/ml streptomycin (Gibco). For gene expression analyses, cells were plated on 35 mm x 10 mm dishes (Corning, USA) or 6-wells plates (Falcon, USA) at a density of 500-1000 neurons/mm^2^. To study the effect of Kdm6 H3K27 demethylase activity inhibition on gene expression, after 2 days *in vitro* (DIV) some cultures were treated with 1.8 μM GSK-J4 (Sigma-Aldrich, USA) or vehicle for 24 h. To study the effect of 17β-estradiol (E2) on *Kdm6a* expression, after 3 DIV the medium of some cultures was replaced for 2 h by fresh medium devoid of B-27 and GlutaMAX-I supplement and then cells were incubated for 2 h with 10^-10^ M E2 (Sigma-Aldrich) or vehicle. For morphometric analysis, cells were plated on 10 mm glass coverslips (Assistent, Germany) at a density of 800 cells/mm^2^. After 3 DIV, some cultures were treated with 1.8 μM GSK-J4 or vehicle for 24 h. The surfaces of glass coverslips and plates were pre-coated with 1 μg/μl poly-L-lysine (Sigma-Aldrich).

### RNA isolation and quantitative Real-Time PCR (qPCR)

Total RNA was extracted from cell cultures and hypothalamic tissue using TRIzol reagent (Invitrogen, USA), purified, and quantified by spectrophotometry on NanoDrop 2000 (Thermo Fisher Scientific, USA) as previously described [41]. 1 μg of RNA per sample was reverse transcribed to cDNA in a 20 μl reaction using M-MLV reverse transcriptase (Promega, USA) and random primers (Invitrogen), following manufacturer’s instructions. qPCR reactions were performed on a StepOne Real Time PCR System using TaqMan or SYBR Green Universal PCR Master Mix (Applied Biosystems, USA) and *Rn18s* (18S rRNA) as control housekeeping gene. TaqMan probes and primers for *Ngn3* were assay-on-demand gene expression products (Applied Biosystems). All other primers (Table 1) were designed using the on-line Primer-Basic Local Alignment Search Tool (Primer-BLAST; National Institutes of Health, USA), selecting primer pairs spanning an exon-exon junction to restrict amplification specifically to mRNA. Primers were verified to amplify with a 95-100% efficiency by performing 4-point calibration curves. Relative quantification of mRNA expression was determined with the ΔΔCt method. Control XYm or control male samples were used as reference group for experiments with FCG or wild-type mice, respectively.

**Table 1.**
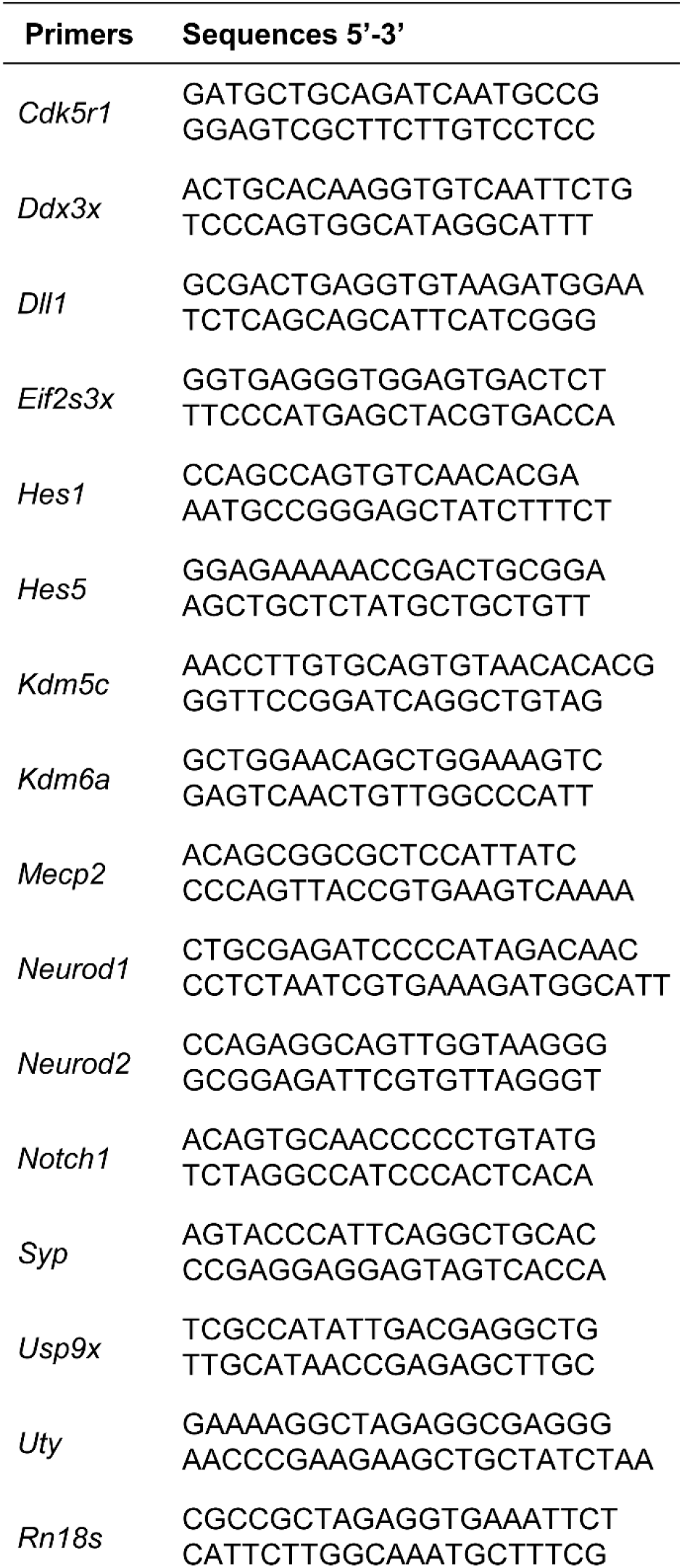
Primer sequences used for gene expression assays by qPCR

### Small interfering RNA (siRNA) transfection

A mixture of 4 different siRNA sequences specific to *Kdm6a* was used at a final concentration of 40 nM total RNA during transfection (1: AGUUAGCAGUGGAACGU-UA, 2: GGACUUGCAGCACGAAUUA, 3: GGUACGGCCUACUGGAAUU, 4: CCACGU-UGGUCAUACUAUA; ON-TARGETplus Mouse Kdm6a Set of 4 siRNA, Dharmacon, UK). A non-targeting siRNA sequence (ntRNA; Dharmacon) was used as control and co-transfection with pmaxGFP (Lonza, Switzerland) was performed in all cases for transfected neurons identification. The efficacy of siRNA targeting *Kdm6a* was assessed by electroporation of neurons using a 4D-Nucleofector X Unit and the corresponding P3 Primary Cell nucleofection kit (Lonza) according to the manufacturer’s instructions, followed by qPCR for *Kdm6a* and *Uty* after 16 h of knockdown. The transfection efficacy by electroporation was 15%, calculated as the percentage of GFP-expressing cells over total number. For morphometric analysis, neurons were transfected at 3 DIV with siRNA targeting *Kdm6a* or ntRNA using Effectene Transfection Reagent (Qiagen, Germany) according to the manufacturer’s instructions. After 16 h of knockdown, cultures were processed for axonal length measurement. The transfection efficacy by this method was 0.1%.

### Western blot

2 DIV hypothalamic neurons were treated with 0.5, 1, 1.8 μM GSK-J4 (Sigma-Aldrich) or vehicle for 24 h and then washed, harvested at 4 °C in RIPA buffer with protease and phosphatase inhibitors, and proteins were resolved and transferred onto nitrocellulose membranes (GE Healthcare, UK) as previously described [44]. Membranes were blocked 90 min at room temperature (RT) in Tris-buffered saline containing 0.1% Tween 20 and 5% BSA, and then incubated overnight at 4 °C with 1:1000 anti-H3K27me3 primary antibody (Cell Signaling Technology, USA). After that, membranes were incubated 1 h at RT with 1:10000 infrared dye-conjugated secondary antibody (LI-COR, USA) and proteins were visualized by Odyssey Infrared Imaging System (LI-COR). After H3K27me3 visualization, blots were stripped and then re-probed with 1:2000 anti-total H3 primary antibody (Cell Signaling Technology) to ensure equal protein loading. Densitometric analyses were performed with the ImageJ software (National Institutes of Health; freely available at https://imagej.nih.gov).

### Immunocytochemistry

After treatment, neurons were fixed for 20 min at RT in 4% paraformaldehyde prewarmed to 37 °C. Transfected neurons were rinsed and mounted on glass slides immediately after, while neurons treated with GSK-J4 were rinsed, permeabilized for 6 min with 0.12% Triton-X plus 0.12% gelatin in phosphate buffered saline (PBS), blocked 1 h at RT in PBS/gelatin, and incubated for 1 h at RT with anti-microtubule associated protein 2 (MAP2) mouse monoclonal antibody (diluted 1:200 in PBS/gelatin; Sigma-Aldrich) and with anti-Tau rabbit polyclonal antibody (diluted 1:500 in PBS/gelatin; Abcam, UK). After rinsing with PBS, cells were incubated for 1 h at RT with the secondary antibodies Alexa 594 goat anti-mouse, for the detection of MAP2, and Alexa 488 goat anti-rabbit, for the detection of Tau (diluted 1:1000 in PBS/gelatin; Jackson ImmunoResearch, USA). Finally, neurons were mounted on glass slides using gerbatol (0.3 g/ml glycerol, 0.13 g/ml Mowiol, 0.2 M Tris-HCl, pH 8.5) plus 1:5000 DAPI for nuclei staining.

### Morphometric analysis

Morphometric evaluation consisted in the assessment of neuritic arborization and axonal length, using in all cases digitized images of the immunostained neurons obtained at 20x magnification with a standard Leica DMI 6000 fluorescence microscope (Leica, Germany) equipped with a digital camera of the same firm. At 4 DIV, the minor processes/dendrites were identified as MAP2 immunoreactive neurites with acute-angled branching, whereas the axon was recognized as a single, thinner neurite of homogeneous caliber along its entire length, at least three times longer than the other processes, with right-angled branching and Tau selective immunoreactivity. Evaluation of neuritic arborization was carried out by a Sholl analysis [45] performed with the CellTarget software ([46]; freely available at https://www3.uah.es/biologia_celular/JPM/CellTarget/CellTarget.html). A grid of 6 concentric circles with increasing radius of 5 μm one respect to the previous one was placed centered on the cell soma, and the number of times neurites intersected any circle was counted in 30 neurons per experimental condition and culture (4 independent cultures). The total number of intersections per neuron was obtained and neuritic arborization complexity was estimated by the Branching Index (BI; [46]). Axonal length was randomly measured in 50-80 neurons per experimental condition and culture (4 independent cultures) using the NeuronJ ImageJ plugin ([47]; freely available at https://imagescience.org/meijering/software/neuronj/).

### Statistical analysis

Data are presented as mean ± SEM and were statistically evaluated by one-, two- or three-way analysis of variance (ANOVA) with treatment, gonadal sex and/or sex chromosome complement as independent variables. ANOVA was followed by *post hoc* comparisons of means by Fisher’s Least Significant Difference (LSD) test. Statistical analysis was performed entirely with Statistica 8 software (StatSoft Inc., USA). p < 0.05 was considered statistically significant. Sample size (n) for all experiments was 3-9 individuals/independent cultures and it is indicated in the figure legends. The number of independent cultures corresponds to the number of pregnant mothers or the number of embryos of each genotype and treatment for wild-type or FCG cultures, respectively. FCG embryos were obtained from 3-5 pregnant mothers per experiment.

## RESULTS

### *Kdm6a*, *Ddx3x*, and *Eif2s3x* expression is higher in XX hypothalamic neurons regardless of gonadal sex

First of all, we studied the expression of several X-linked genes particularly interesting for their involvement in neuronal growth and differentiation, in order to identify those with higher expression levels in XX than in XY neurons. Primary cultures of hypothalamic neurons from E14 FCG mice were established, maintained during 3 DIV, and then processed to determine the relative mRNA levels of seven X-linked genes by qPCR: *Kdm6a*, *Eif2s3x*, *Ddx3x*, *Kdm5c*, *Mecp2*, *Usp9x*, and *Syp* (Fig. 1). *Kdm6a*, *Ddx3x* and *Eif2s3x* showed significantly higher expression levels in hypothalamic neurons carrying the XX sex chromosome complement compared to those carrying the XY, regardless of the gonadal sex of the donor embryos (XXf and XXm > XYf and XYm; ANOVA *Kdm6a*: F(1, 18) = 6. 58, p = 0.0194; ANOVA *Ddx3x*: F(1,19) = 14.14, p = 0.0013; ANOVA *Eif2s3x*: F(1,17) = 5.79, p = 0.0278). On the other hand, no differences by either sex chromosome complement or gonadal sex were found in *Kdm5c*, *Mecp2*, *Usp9x*, and *Syp* mRNA levels.

**Fig. 1.**
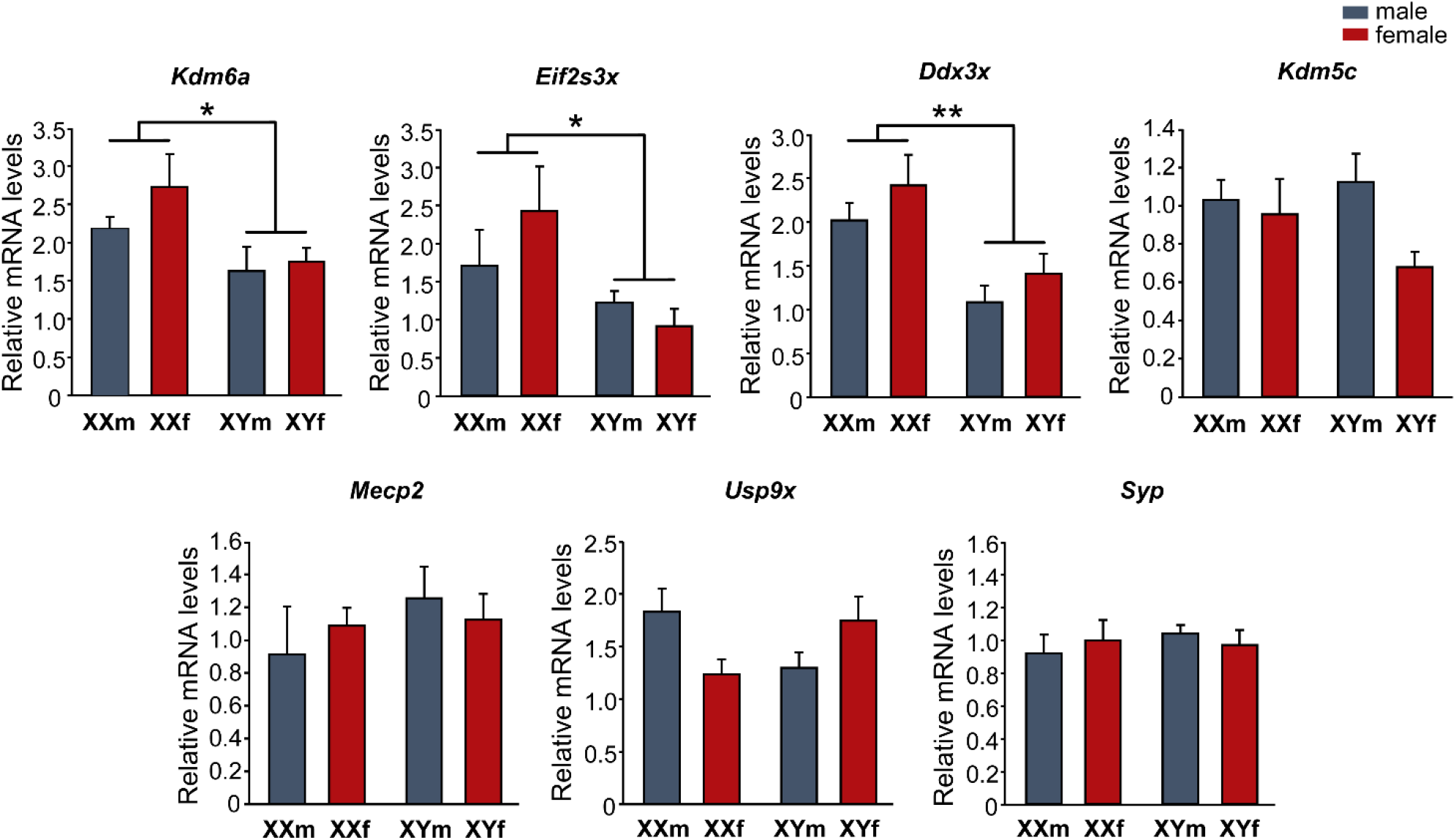
Relative mRNA levels of X-linked genes *Kdm6a*, *Eif2s3x*, *Ddx3x*, *Kdm5c*, *Mecp2*, *Usp9x*, and *Syp* in FCG hypothalamic neurons at 3 DIV. *Kdm6a*, *Eif2s3x*, and *Ddx3x* show higher expression levels in XX compared to XY cultures, regardless of gonadal sex (XXf and XXm > XYf and XYm). Data are mean ± SEM. n = 5-7 independent cultures for each genotype. \**p* < 0.05; \*\**p* < 0.01

### Higher *Kdm6a* expression levels in XX neurons do not change with E2 treatment or age

Given the role of *Kdm6a* as a key epigenetic regulator of gene transcription, and considering the results showing its sexually dimorphic expression determined by the sex chromosome complement (XX > XY), we decided to focus our study henceforth on its involvement in the sex-specific differentiation of hypothalamic neurons. First, knowing the importance of E2 as the main sex hormone mediating organizational processes in the rodent brain, we treated cultures with this estrogen and analyzed the effect on *Kdm6a* mRNA levels. Remarkably, E2 presence in the culture milieu did not alter *Kdm6a* gene expression levels, maintaining higher values in XX than in XY neurons, independently of sex and treatment (Fig. 2; ANOVA: F(1, 28) = 14.14, p = 0.00079).

**Fig. 2.**
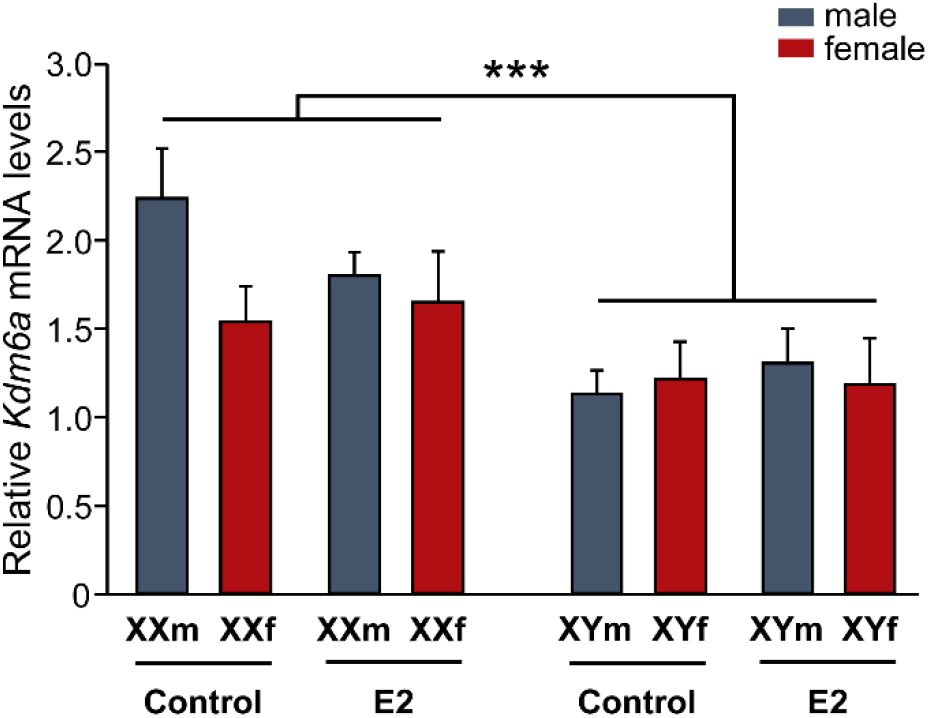
17β-estradiol (E2) effect on *Kdm6a* mRNA levels in FCG hypothalamic neurons. Incubation with 10^-10^ M E2 did not change *Kdm6a* gene expression levels, remaining higher in XX vs. XY cultures, regardless of sex and treatment. Data are mean ± SEM. n = 4-5 independent cultures for each genotype and treatment. \*\*\**p* < 0.001

Moreover, *Kdm6a* expression pattern did not change either with age, remaining higher in the hypothalamic tissue of XX than XY animals at E14, postnatal day 0 (P0, newborns), and postnatal day 60 (P60, gonadally intact adults), regardless of whether individuals carried testes or ovaries (Fig. 3; ANOVA E14: F(1,19) = 87.74, p = 0.000000015; ANOVA P0: F(1,13) = 12.49, p = 0.0037; ANOVA P60: F(1,15) = 11.15, p = 0.0045).

**Fig. 3.**
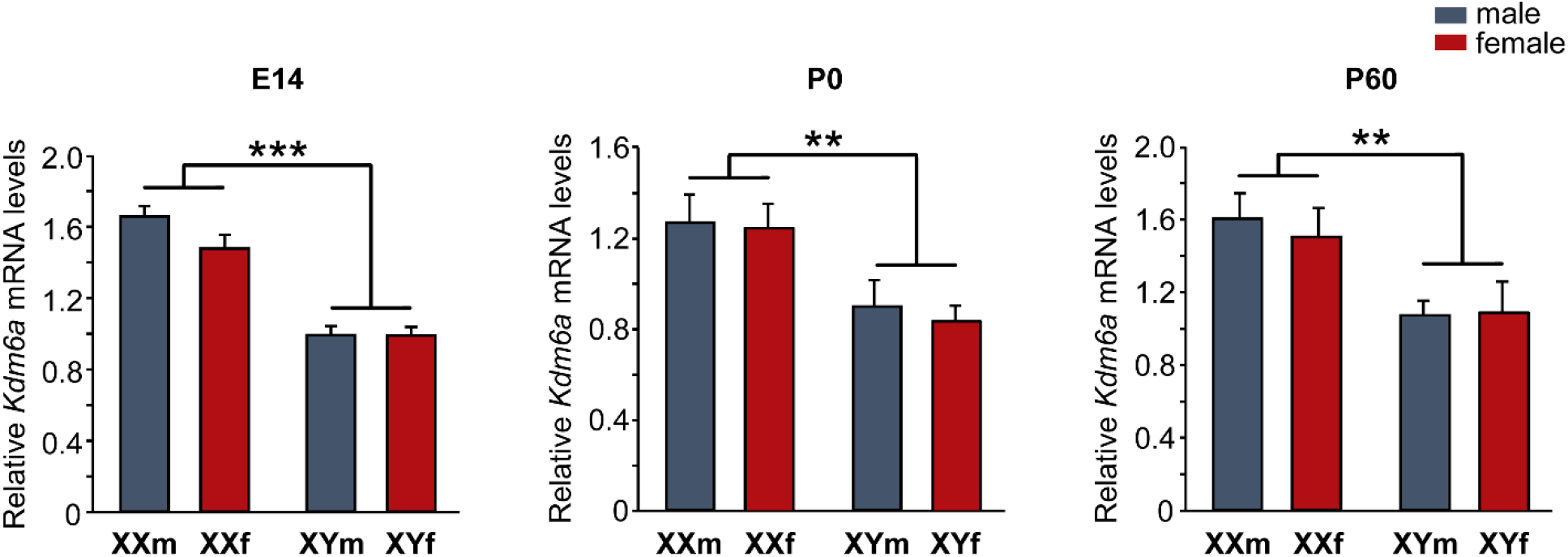
Relative *Kdm6a* mRNA levels in hypothalamic tissue of FCG mice at E14, P0, and P60. Gene expression was higher in XX than XY animals at all ages and independently of gonadal sex. Data are mean ± SEM. n = 5-6 individuals for each genotype and age. \*\**p* < 0.01; \*\*\**p* < 0.001

### Kdm6 H3K27 demethylase activity and higher *Kdm6a* gene expression are required for the differentiation of longer axons in XX hypothalamic neurons

In order to analyze the requirement for Kdm6 H3K27 demethylase activity in the sexually dimorphic differentiation of hypothalamic neurons, we employed the specific cell permeable small-molecule inhibitor GSK-J4 [48]. First, we validated the effectiveness of GSK-J4 blocking Kdm6 demethylase activity in primary cultures of hypothalamic neurons from wild-type CD1 mice, performing a dose/response curve and evaluating the H3K27me3 levels by Western blot. In agreement with times and concentrations previously reported by others [49], treatment of cultures for 24 h with GSK-J4 1.8 μM showed the highest H3K27me3 levels (Fig. 4a) without affecting cell viability, proving to be effective in inhibiting the Kdm6 enzymatic activity. Besides, treatment did not affect the gene expression levels of *Kdm6a* or its paralogue on the Y chromosome, *Uty*, presenting neuronal cultures derived from female embryos higher *Kdm6a* mRNA levels than those from male, both in control and treated conditions (Fig. 4b; ANOVA: F(1, 13) = 10.25, p = 0.007). Considering these results, we selected this condition for all further experiments using GSK-J4 to inhibit Kdm6 demethylase activity.

**Fig. 4.**
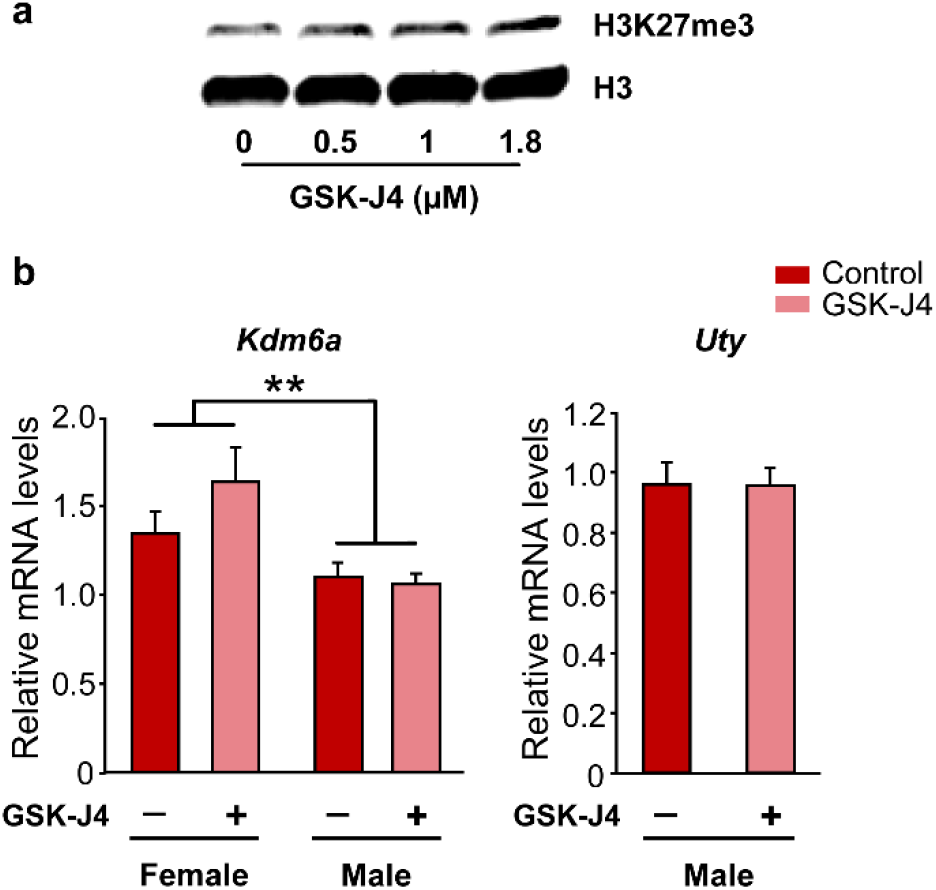
Validation of GSK-J4 treatment as an inhibitor of Kdm6 demethylase activity in wild-type hypothalamic neurons. **a** Representative immunoblot showing the increase in histone 3 lysine 27 trimethylation (H3K27me3) levels as a result of Kdm6 demethylation blockade for 24 h at increasing concentrations of GSK-J4. Total histone 3 (H3) was used as loading control. **b** GSK-J4 1.8 μM treatment for 24 h inhibited demethylation without affecting *Kdm6a* or *Uty* gene expression. *Kdm6a* mRNA levels were higher in female than male cultures, regardless treatment. Data are mean ± SEM. n = 3-6 independent cultures for each sex and treatment. \*\**p* < 0.01

Thereafter, we evaluated the effect of Kdm6 H3K27 demethylase activity inhibition by GSK-J4 over neuronal growth and differentiation *in vitro*, analyzing neuritic arborization complexity and axonal length as parameters of neuronal morphology. Assessment of neuritic arborization was performed by a Sholl analysis, counting per cell the number of times neurites intersected any line of the concentric circle grid (Fig. 5a). At this level, significant differences were only observed in the number of intersections due to sex, with no effect of treatment: neurons derived from female embryos presented a higher mean number of intersections than those from male embryos, independently of GSK-J4 blockade (Fig. 5b; ANOVA: F(1, 12) = 5.15, p = 0.0425). Accordingly, calculation of the BI estimated a higher complexity in the neuritic arborization of female (BI = 14.91 ± 0.85) than male (BI = 11.26 ± 1.25) neurons, without treatment effect (ANOVA: F(1, 12) = 6.067, p = 0.03). Regarding axonal length analysis, in agreement with results previously reported by our laboratory [34, 35], a clear sexual dimorphism was observed in control conditions, showing female neurons significantly longer axons on average than male neurons. Remarkably, GSK-J4 treatment nullified this difference, decreasing axonal length only in female derived neurons (Fig. 5c and d; ANOVA: F(1, 12) = 22.86, p = 0.0004).

**Fig. 5.**
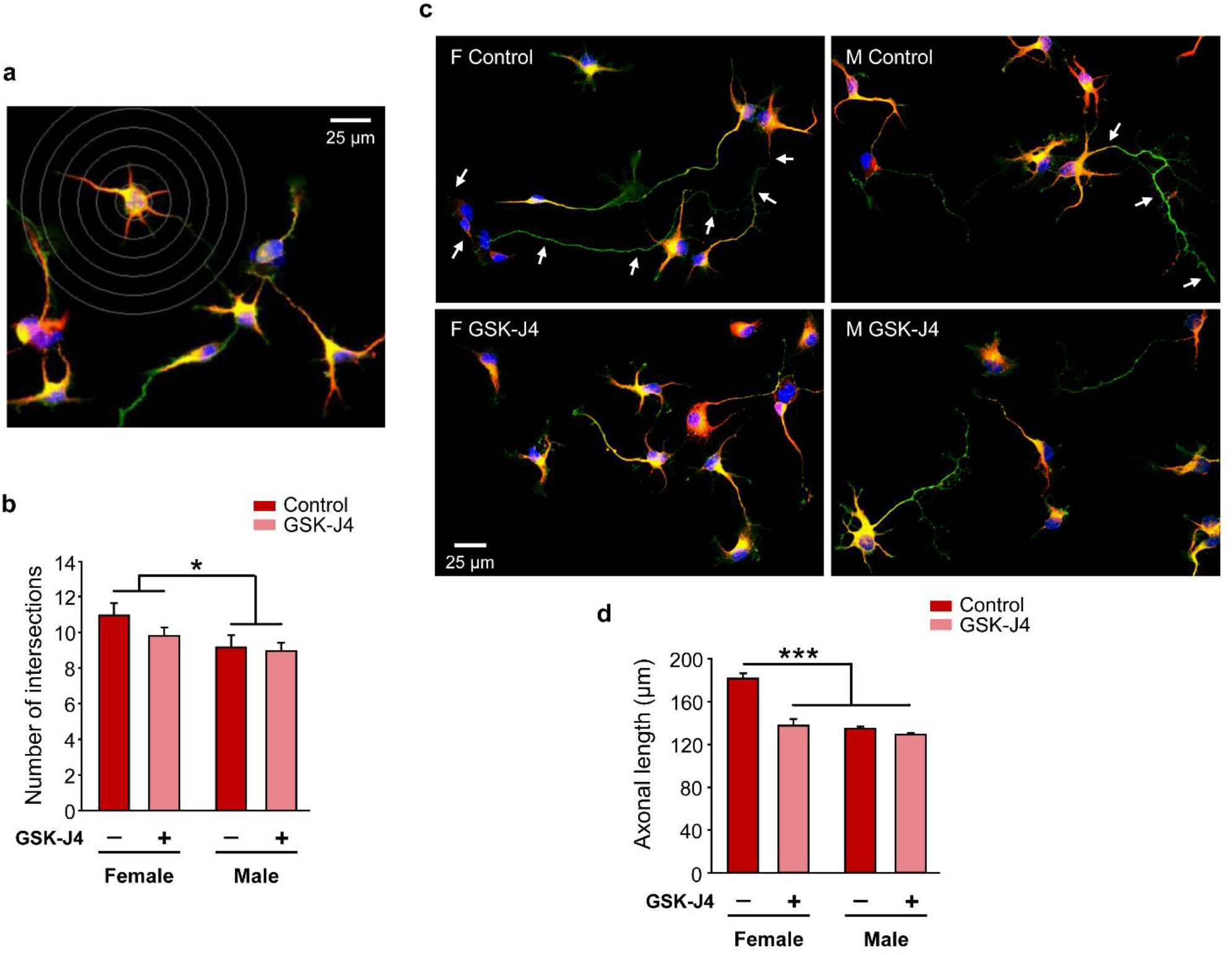
Sexually dimorphic neuritogenesis of hypothalamic neurons depends on Kdm6 H3K27 demethylase activity. **a** Representative fluorescence image of female control neurons showing the grid used for Sholl analysis. **b** Number of intersections of neurites with Sholl’s grid of female and male neurons treated (GSK-J4) or not (Control) at 3 DIV with GSK-J4 1.8 μM for 24 h. Female neurons showed higher values than male neurons, irrespective of treatment. **c** Representative fluorescence images of female (F) and male (M) neurons treated (GSK-J4) or not (Control) at 3 DIV with GSK-J4 1.8 μM for 24 h. Arrows indicate the axon of a female control neuron and a male control neuron along their full length. **d** Mean axonal length for each experimental condition. GSK-J4 treatment abolished sex differences decreasing axonal length only in female neurons. Data are mean ± SEM. n = 4 independent cultures for each sex and treatment. \**p* < 0.05; \*\*\**p* < 0.001

Given these results that suggest a participation of Kdm6 enzymes in the sexually dimorphic regulation of axogenesis in hypothalamic neurons, and that treatment with GSK-J4 inhibits the activity of both Kdm6a and Kdm6b, being impossible by this methodological approach to discern the individual contribution of each demethylase, we proceeded to the specific knockdown of *Kdm6a* transcripts using siRNA designed to downregulate the expression of this gene without affecting *Kdm6b* or *Uty*. Once the effectiveness of siRNA decreasing *Kdm6a* mRNA levels was proven (Fig. 6b; (ANOVA: F(1, 7) = 6.06, p = 0.043), neuronal cultures derived from sex-segregated wild-type mice were transfected to study the effect of *Kdm6a* knockdown on axonal growth. Again, as in the results obtained using GSK-J4, morphometric analysis showed a significant decrease in axonal elongation in female but not in male derived neurons after *Kdm6a* downregulation, abolishing the sex differences observed in axonal length in control conditions (Fig. 6a and c; ANOVA: F(1, 9) = 6.16, p = 0.0348).

**Fig. 6.**
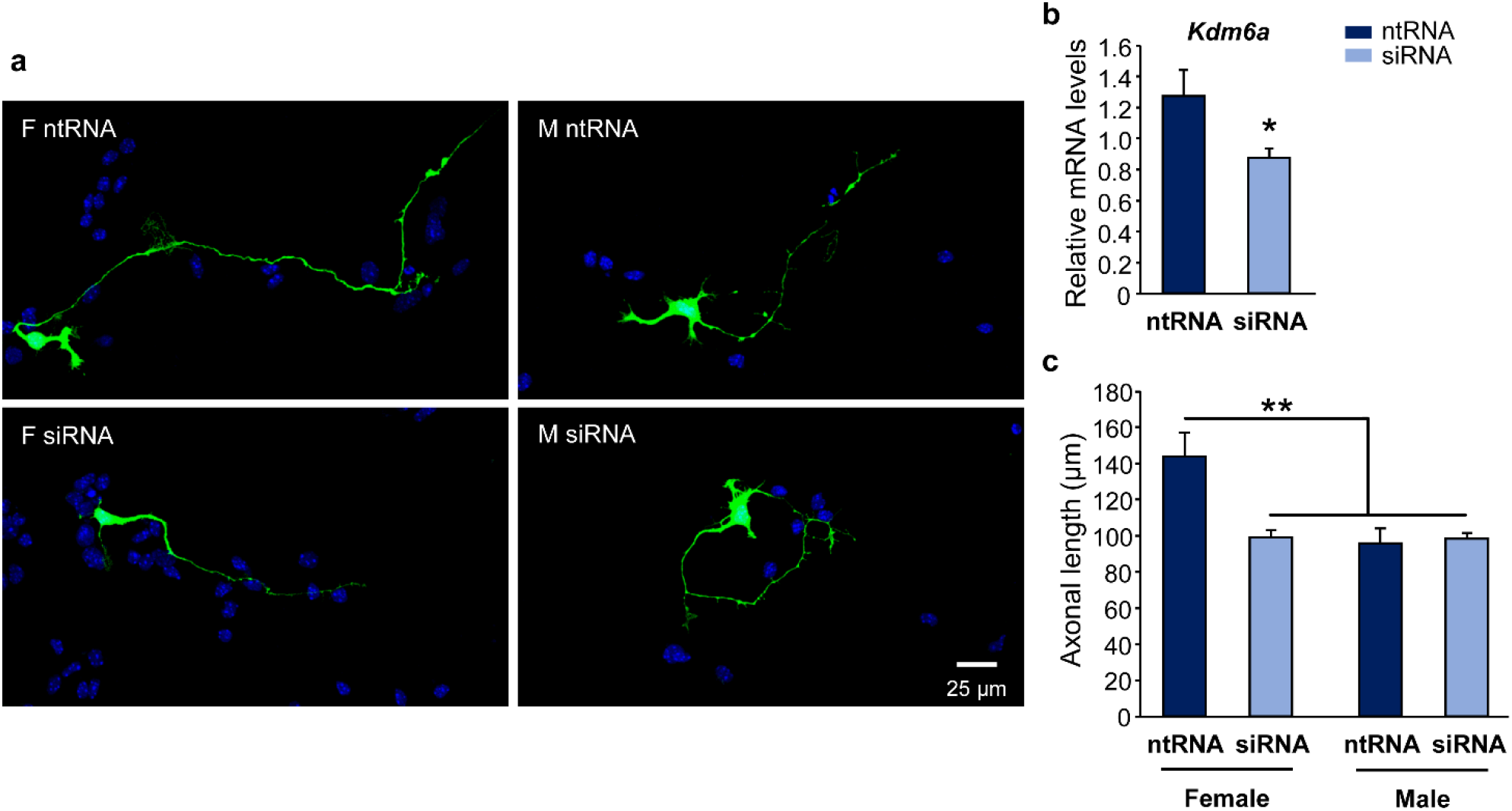
Higher *Kdm6a* gene expression is required for the differentiation of longer axons in female hypothalamic neurons. **a** Representative fluorescence images of female (F) and male (M) neurons co-transfected with a non-targeting siRNA sequence (ntRNA) and GFP or siRNA targeting *Kdm6a* (siRNA) and GFP at 3 DIV for 16 h. **b** Targeting siRNA were effective downregulating *Kdm6a* mRNA levels in hypothalamic neurons *in vitro*. **c** Mean axonal length for each experimental condition. *Kdm6a* knockdown abolished sex differences decreasing axonal length only in female neurons. Data are mean ± SEM. n = 4 independent cultures for each sex and treatment. \**p* < 0.05; \*\**p* < 0.01

### Kdm6 H3K27 demethylase activity regulates the expression of *Ngn3* in a sex-specific manner determined by the sex chromosome complement

Since in previous work we have demonstrated that higher expression of *Ngn3* in XX hypothalamic neurons is required for faster maturation and longer axons of these neurons *in vitro* [34], we investigated whether *Kdm6a* regulates the expression of *Ngn3* and other neuritogenic genes such as *Neurod1*, *Neurod2*, and *Cdk5r1*. Cultures from E14 wild-type mice segregated by sex were maintained 2 DIV and then treated with GSK-J4 for 24 h before processing for gene expression analysis by qPCR. As in Scerbo *et al*. [34], neurons derived from female embryos presented higher *Ngn3* mRNA levels than those from males under control conditions (Fig. 7a). Interestingly, inhibition of Kdm6 demethylase activity significantly reduced *Ngn3* expression in female cultures, while increasing it in males (ANOVA: F(1, 24) = 14.75, p = 0.0008). Moreover, GSK-J4 treatment significantly reduced *Neurod1*, *Neurod2*, and *Cdk5r1* gene expression equally in both sexes (Fig. 7a; ANOVA *Neurod1*: F(1, 25) = 13.5, p = 0.001; ANOVA *Neurod2*: F(1, 24) = 10.15, p = 0.0039; ANOVA *Cdk5r1*: F(1, 18) = 4.41, p = 0.049).

**Fig. 7.**
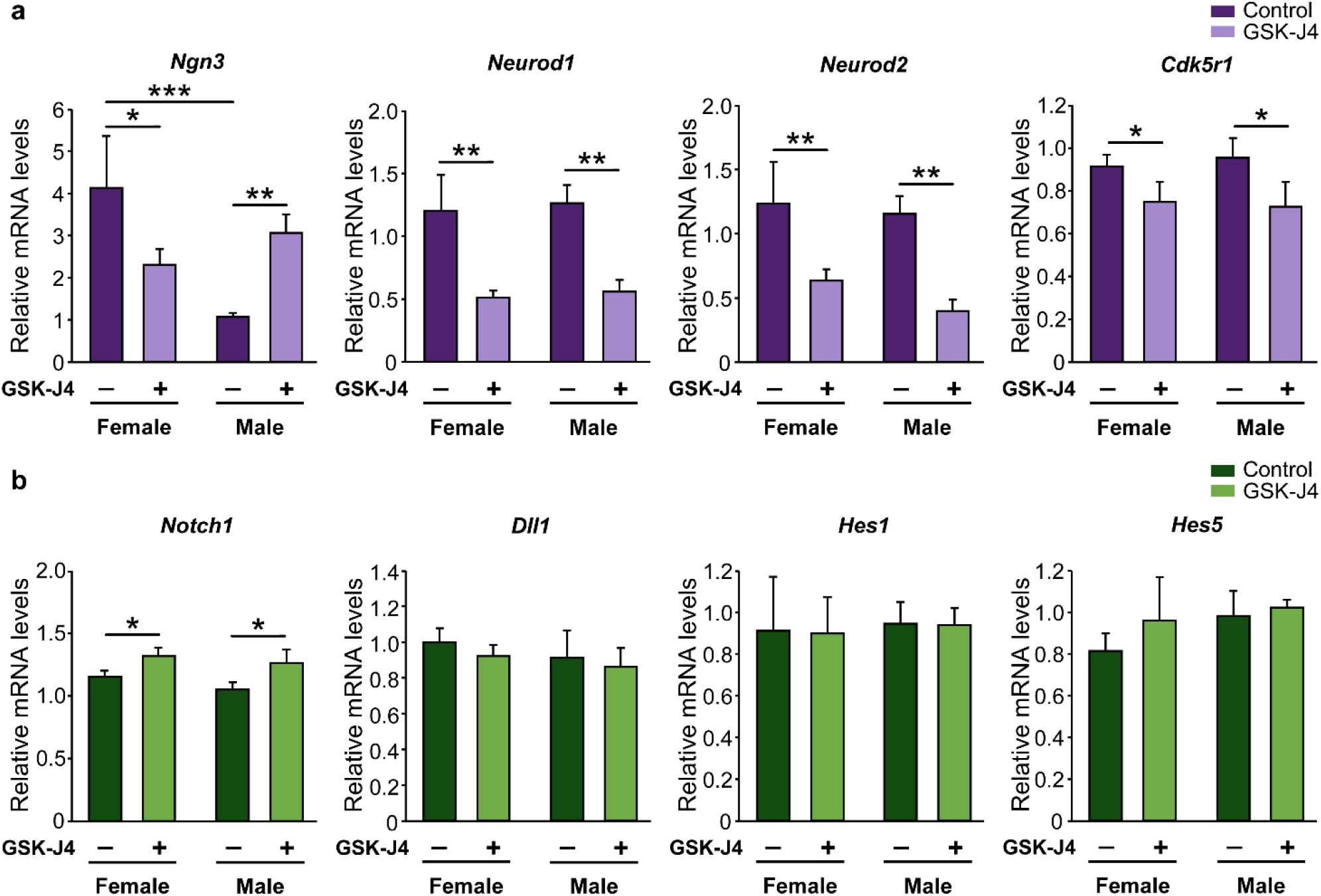
Effect of Kdm6 H3K27 demethylase activity inhibition on neuritogenesis-related genes expression in wild-type hypothalamic neurons. **a** Effect of GSK-J4 treatment on neuritogenic genes expression. Kdm6 demethylases blockade reduced *Neurod1*, *Neurod2* and *Cdk5r1* mRNA levels in both sexes equally, while a sex-specific effect was observed on *Ngn3* levels, with a decrease in female and an increase in male neurons. **b** Effect of GSK-J4 treatment on Notch signaling genes expression. Kdm6 demethylases blockade did not change *Dll1*, *Hes1*, and *Hes5* mRNA levels, while *Notch1* expression increased with treatment both in female and male neurons. Data are mean ± SEM. n = 4-9 independent cultures for each sex and treatment. \**p* < 0.05; \*\**p* < 0.01; \*\*\**p* < 0.001

Next, we evaluated the effect of the Kdm6 demethylase activity inhibition over the expression of some of the main Notch signaling elements, knowing that *Ngn3* transcription is downregulated when this pathway is activated [50, 51]. Statistical analysis showed no significant differences due to gonadal sex or GSK-J4 treatment in *Dll1*, *Hes1*, and *Hes5* relative mRNA levels, while *Notch1* expression increased with treatment both in female and male neurons (Fig. 7b; ANOVA *Notch1*: F(1, 18) = 5.68, p = 0.028).

Finally, neuronal cultures derived from FCG transgenic mice were treated with the GSK-J4 inhibitor and processed for *Ngn3* relative expression analysis. In agreement with our previous results [34, 35], sexually dimorphic expression of *Ngn3* was proven to be determined by sex chromosome complement in control conditions, with XX neurons expressing significantly higher levels than XY neurons regardless of gonadal sex. Remarkably, the inhibition of Kdm6 H3K27 demethylase activity in FCG transgenic cultures showed sex chromosomes-specific effects in line with the results observed in wild-type cultures: a decrease in *Ngn3* mRNA levels in XX neurons and an increase in XY neurons due to GSK-J4 treatment and irrespective of gonadal type (Fig. 8; ANOVA: F(1, 21) = 29.646, p = 0.00002).

**Fig. 8.**
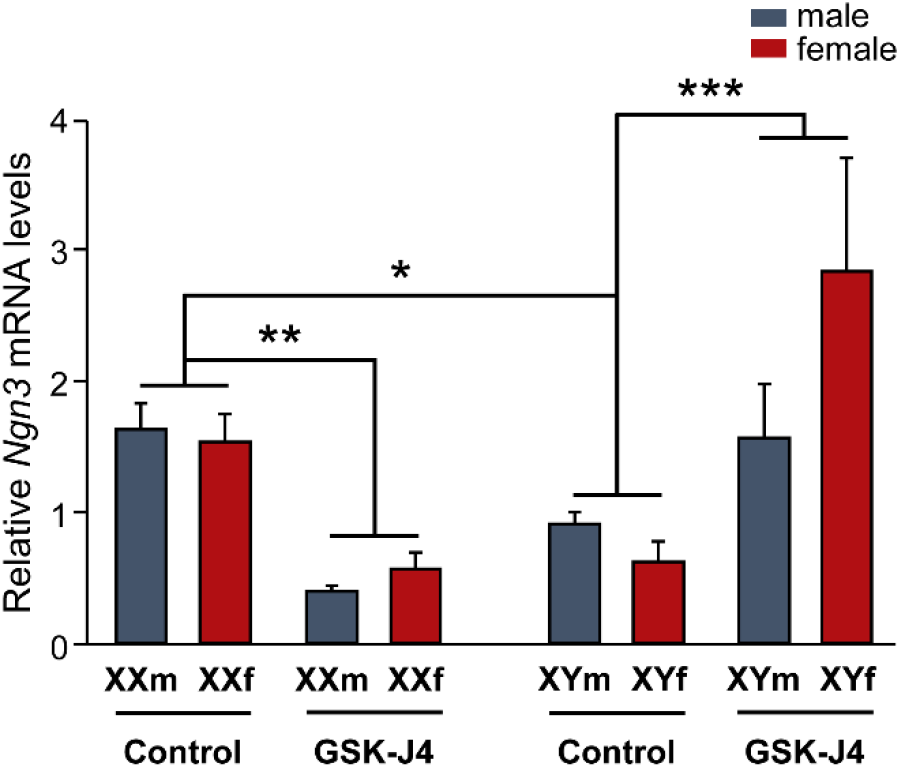
Kdm6 H3K27 demethylation regulates *Ngn3* expression in a sex-specific manner determined by the sex chromosome complement. In control conditions, FCG XX hypothalamic neurons express higher *Ngn3* mRNA levels than XY neurons, regardless of gonadal sex. Kdm6 demethylases blockade by GSK-J4 generated a decrease in *Ngn3* mRNA levels in XX and an increase in XY neurons. Data are mean ± SEM. n = 3-5 independent cultures for each genotype and treatment. \**p* < 0.05; \*\**p* < 0.01; \*\*\**p* < 0.001

## DISCUSSION

Currently it is well known that the sex chromosome complement contribution to brain sexual differentiation is not just limited to the determination of a gonadal type *in utero* (testes for XY, ovaries for XX), but continues playing a role in the development and expression of sex dimorphisms at morphological, physiological, and behavioral levels throughout life. However, little is known about the identity of particular X or Y-linked genes involved and the mechanisms by which these genes participate in the process. Here we provide valuable evidence pointing to *Kdm6a* as one of the main X-linked mediators of the sexually dimorphic differentiation of hypothalamic neurons, regulating sex differences in the expression of the proneural gene *Ngn3* and axonal growth.

Although the X chromosome comprises only about 5% of the mouse and human genomes, it is particularly enriched in brain relevant genes, containing five times as many genes linked to the development and physiology of the nervous system compared to an autosome [6, 52–54]. Many of these genes escape XCI, although expression levels from the Xi vary between escapees and can be tissue-, developmental stage-, species-, individual-, and even single cell-dependent for the same gene [15, 55, 56]. Among the seven X-linked genes whose relative expression we analyzed by qPCR, all show previous evidence of XCI escape in different tissues in both mice and humans with the exception of *Mecp2* [57–62]. Our results show that while *Kdm6a*, *Eif2s3x*, and *Ddx3x* presented higher expression levels in XX compared to XY hypothalamic neurons and independently of gonadal sex, *Kdm5c*, *Mecp2*, *Usp9x*, and *Syp* did not vary in expression among the four genotypes in the FCG model. In the mouse nervous system, *Kdm6a*, *Kdm5c*, *Eif2s3x*, and *Ddx3x* have been shown to escape XCI in studies using whole brain [14, 59, 63, 64], while few studies have assessed this question specifically by brain region and cell type. Armoskus *et al*. [65] demonstrated, by mRNA extraction from tissue blocks encompassing cerebral cortex and hippocampus of neonatal mice, a higher expression of *Kdm6a* and *Eif2s3x* in females compared to males. On the other hand, Xu *et al*. [66] have reported higher levels of *Kdm6a* mRNA in adult females in cortex, striatum, suprachiasmatic nucleus, ventromedial hypothalamus, hippocampus (CA1/CA3), dentate gyrus, and habenula. In turn, the same study demonstrated that this dimorphism is sex chromosome complement dependent, as higher *Kdm6a* expression was observed in whole brain of adult XX FCG mice (both gonadal males and females) compared to XY mice. As far as we are aware, the present is the first study to explore and demonstrate the sexually dimorphic expression of *Kdm6a*, *Eif2s3x*, and *Ddx3x* specifically in hypothalamic neurons prior to the organizational action of gonadal hormones. The use of the FCG transgenic model and the election of a developmental time prior to the critical period of brain hormonal organization allows us to present robust results clearly showing that the sexually dimorphic expression of *Kdm6a*, *Eif2s3x*, and *Ddx3x* is determined by the sex chromosome complement (XX > XY), and strongly suggesting that these genes escape XCI specifically in hypothalamic neurons during early development.

Considering the sex differences in *Kdm6a*, *Eif2s3x*, and *Ddx3x* expression and their regulatory functions either at the transcriptional or translational level, it is possible to hypothesize that these genes could be involved in the definition of sexual dimorphisms during brain development. *Ddx3x* encodes for an evolutionarily conserved DEAD-box RNA helicase involved in a multiplicity of fundamental cellular processes, including transcription, splicing, mRNA transport, translation, and regulation of the Wnt/β-catenin signaling pathway [67–69]; mutations in this gene have been reported in different types of cancer and associated with intellectual disability [70]. On the other hand, little is known about the functions of *Eif2s3x* beyond the fact that it is involved in the regulation of protein synthesis as a subunit of the translation initiation factor eIF2 [71, 72]. Finally, particularly interesting is *Kdm6a*, which protein product plays important roles as a genome-wide regulator of gene expression not only by removing H3K27me2/me3 repressive marks but also by directly interacting with promoter regions [19, 20]. Thus, considering the key role of *Kdm6a* as an epigenetic regulator of transcription and its higher expression in XX hypothalamic neurons, we set out to further study its participation in the sexually dimorphic differentiation of these neurons.

In rodents, E2 converted from testosterone by the enzyme P-450 aromatase is by far the main steroid hormone responsible for brain masculinization and defeminization during the so-called critical or sensitive period of sexual differentiation, which comprises the E17-P10 perinatal phase of development [42, 73]. Surprisingly, no significant differences in *Kdm6a* mRNA levels were found between E2-treated and unstimulated control cultures, always observing higher *Kdm6a* levels in XX neurons, irrespective of sex and treatment. Many estrogen actions depend on mediation or interaction with other trophic factors, such as neurotrophins (BDNF, IGF-I, GDNF), neurotransmitters (GABA, glutamate), and many other regulatory molecules (prostaglandins, neuroglobin, huntingtin, etc.), often produced and secreted by cells other than neurons, as is the case with glial cells in nervous tissue [73–78]. For such reason we decided to validate *in vivo* what was observed in primary cultures, evaluating in tissue from the hypothalamic region whether the expression pattern of *Kdm6a* changes depending on the sexually dimorphic fluctuations in circulating levels of gonadal hormones that occur throughout life. Consistent with the results from *in vitro* stimulation with E2, the higher *Kdm6a* expression in XX individuals and independent of gonadal sex remained unaltered in the hypothalamus at E15, P0, and P60 (before, during, and after critical period, respectively), reinforcing evidence that the sex-specific expression pattern of the demethylase is directly determined by the number of X chromosomes present in each cell and would not be modulated by gonadal hormones.

Our laboratory has previously demonstrated and reported a sexual dimorphism in hypothalamic neurons differentiation prior to the action of gonadal hormones and dependent on sex chromosome complement during brain development: XX neurons show an accelerated neuritic development *in vitro* compared to XY neurons, in terms of axonal length and dendrite development and branching [34, 35]. For example, at 2 DIV cultures, it was observed that while 14% of the female neurons had reached stage III of *in vitro* development (polarized neurons), none of the male neurons had developed axon yet. In addition, female cultures showed a higher proportion of cells with branched neurites than male cultures. Similar conditions with female neurons showing greater neuritic development compared to male neurons were observed from 1 to 5 DIV, but no longer at 6 and 7 DIV [34]. Interestingly, these sexual differences disappear under E2 stimulation, as only XY neurons respond to the estrogen by accelerating their development, elongating their axons in a clear axogenic effect, and matching XX neurons, suggesting that a similar effect could occur *in vivo* with the arrival of early gonadal secretions and their organizing actions on brain during perinatal male development [34, 35, 78, 79]. Reisert *et al*. [80] and Ruiz-Palmero *et al*. [81] have reported similar results by observing that neuritic development initially occurs faster in female than in male cultures of rat diencephalon dopaminergic neurons and mouse hippocampal neurons, respectively. As discussed in Scerbo *et al*. [34], although the biological meaning of these transient sex differences in neuronal development is unknown, it is conceivable that such differences could lead *in vivo* to the establishment of sexually dimorphic synaptic connectivity patterns: female neurons, by differentiating first, could find a greater number of available presynaptic sites when reaching target regions and/or present more free postsynaptic sites when receiving early synaptic afferents. Thus, through these mechanisms, female and male hypothalamic neurons could finally establish different connectivity patterns.

The results of the present study replicate what was previously observed in terms of a greater neuritic development of female hypothalamic neurons and show, in turn, that both the silencing of *Kdm6a* gene expression and the pharmacological blockade of its enzymatic activity affected axonal elongation only in female cultures, generating a decrease in axonal length that was not observed in male cultures. The GSK-J4 prodrug is hydrolyzed inside the cell to GSK-J1, this molecule being the active drug that inhibits H3K27 demethylase activity of the enzyme through competition for the 2-oxoglutarate co-substrate binding site [48]. GSK-J1 has been shown to be able to block the active site and inhibit the demethylase activity of all enzymes of the Kdm6 subfamily, i.e., Kdm6a, Kdm6b and Uty/Kdm6c (although *in vivo* demethylation capacity of the latter is under discussion; [82, 83]). Therefore, the results derived from GSK-J4 treatment of neuronal cultures could be reflecting the effects not only of blocking the demethylase activity of Kdm6a, but also that of the other Kdm6 enzymes. However, the same results were obtained when silencing *Kdm6a* gene expression by siRNA sequences designed to specifically degrade the transcripts of this demethylase without affecting *Kdm6b* and *Uty*. Taken together, the results obtained from both pharmacological blockade by GSK-J4 and *Kdm6a* silencing indicate that early sex differences in axogenesis of hypothalamic neurons depend in part on the sexually dimorphic expression of *Kdm6a*: its H3K27 demethylase activity and increased transcription in XX neurons are necessary in mediating the processes leading to the proper development of these cells, whereas they are not indispensable (at least at this time) for XY neurons. These observations become even more relevant considering previous work by other groups in which, for example, it was shown that while *Kdm6a* knockout (KO) is fatal in female mice at E11-12 (with severe heart malformations and defects in neural tube closure), KO males (albeit at a lower frequency than expected) survive to adulthood and are fertile [82, 84].

Previous studies have demonstrated a requirement for *Kdm6a* during neuronal development, for example, in the determination and differentiation of cortical neurons [23] and in the dendritic development, synapse formation and maturation, and synaptic transmission of mouse hippocampal neurons [27] and human neurons [28]. However, the precise mechanisms through which *Kdm6a* participates in these processes are unknown, although it is assumed to occur through the control of the expression profile of numerous genes directly involved in such processes. Several studies have demonstrated large-scale changes in the transcriptome following deletion or blockade of *Kdm6a*, observing a significant decrease in the expression of genes linked to neurodevelopment, neurogenesis, synaptogenesis, and neurotransmitter transport and secretion, among other relevant neuronal processes [24, 27, 28, 85]. In Scerbo *et al*. [34], our laboratory demonstrated that, prior to the critical period, hypothalamic neurons derived from female embryos exhibit higher levels of the neuritogenic factor *Ngn3* than male neurons. In turn, it was observed that the higher expression of *Ngn3* is necessary for the more accelerated development shown by female hypothalamic neurons *in vitro*. On the other hand, although it was shown in cultures derived from FCG transgenic mice that sexually dimorphic expression of *Ngn3* is determined by the sex chromosome complement, the precise factors and mechanisms by which the X and Y chromosomes would regulate the expression of this autosomal gene are still unknown [34, 35]. Thus, knowing that both *Kdm6a* and *Ngn3* participate in the regulation of sexually dimorphic neuritogenic development of hypothalamic neurons, that both genes show similar expression patterns with higher levels in XX than in XY neurons, and that *Ngn3* expression is determined by the sex chromosome complement, we wondered whether *Kdm6a* would be responsible, at least in part, for controlling the sexually dimorphic expression of this proneural factor. The inhibition of H3K27 demethylation activity of Kdm6 by GSK-J4 generated similar results in cultures from wild-type and FCG transgenic animals in terms of *Ngn3* expression: drug treatment decreased *Ngn3* levels in XX hypothalamic neurons, whereas increasing them in XY neurons. These results suggest that *Ngn3* expression is mediated by H3K27 demethylation, and indicate different regulatory mechanisms for males and females that are sex chromosome complement-dependent and gonadal sex-independent. Since Notch signaling pathway activation inhibits neurogenesis through repression of numerous proneural genes including *Ngn3* [50, 51], we decided to evaluate the effect of blocking H3K27 demethylation on gene expression of some of the main elements of this pathway. Results were equal in both sexes and showed an increase of *Notch1* after GSK-J4 inhibition and no treatment effect on *Dll1*, *Hes1*, and *Hes5* levels, suggesting that the sexually dimorphic regulation of Kdm6 demethylase activity on *Ngn3* would not be mediated by the Notch pathway in hypothalamic neurons. Further experiments are required to completely understand why *Ngn3* is differentially regulated in females and males by H3K27 demethylation.

Taken together, our results point to *Kdm6a* as an X-linked gene directly involved in the sexually dimorphic differentiation of XX hypothalamic neurons, expressing at higher levels in these cells independently of gonadal sex, promoting through its H3K27 demethylase activity the transcription of *Ngn3* and other proneural factors such as *Neurod1*, *Neurod2*, and *Cdk5r1*, and mediating the process of axogenesis in a sex-specific manner prior to hormonal organization of the brain. We have shown that the sex difference in *Kdm6a* transcriptional levels is determined by the sex chromosome complement, while gene expression results from E2-stimulated neuronal cultures and hypothalamus of neonatal and adult FCG animals strongly suggest that *Kdm6a* expression is not subject to sex hormone regulation *per se*. However, we cannot exclude the possibility that the sex-specific effects determined by *Kdm6a* on *Ngn3* expression and axogenesis are mediated by sex steroids, for example, if they occur through a differential regulation of the demethylase on neurosteroidogenesis in XX and XY individuals. In this regard, it would be particularly interesting to develop further experiments focused on the possible involvement of *Kdm6a* in the regulation of the neurosteroidogenic pathway and the generation of sexual dimorphisms in the hypothalamus and other brain regions by this way.

Considering the results of the present study and our own and other groups’ previous evidence, we propose the following hypothesis to explain the factors and mechanisms currently known to act in the sexual differentiation of hypothalamic neurons (Fig. 9). On the one hand, XX neurons, whose embryonic and perinatal development occurs in the absence of significant levels of gonadal hormones, show higher expression of X-linked genes that escape XCI, such as *Kdm6a*, *Ddx3x*, and *Eif2s3x*. Greater *Kdm6a* levels (and possibly other X-linked genes) during early neurodevelopment in XX neurons lead to increased expression of neuritogenic genes such as *Ngn3*, which, in turn, promotes neuritogenesis. On the other hand, XY neurons exhibit lower expression of neuritogenic-related escapee genes. However, these cells are influenced by E2 produced intraneuronally from testosterone secreted by testes during the critical period of perinatal development, presenting, in addition, a higher expression and enzymatic activity of aromatase [86, 87] and a higher sensitivity to E2 than XX neurons by expressing more estrogen receptors *Esr1* and *Esr2* [35, 88]. E2 organizational actions during the critical period in XY neurons would ensure the increase in *Ngn3* expression necessary for the promotion of neuritogenesis. Therefore, in terms of the neuritic development of hypothalamic neurons, different mechanisms would exist depending on the sex chromosome complement to ultimately determine the adequate neuritic arborization and axonal length in both sexes. Thus, dimorphic factors such as, for example, the higher expression of X-linked genes in XX neurons or the early influence of gonadal hormones in XY neurons are compensated or balanced. In this way, starting from a different genetic background, through sexually dimorphic mechanisms, a similar neuritic development is reached in both sexes. Following this line of reasoning, evolution would have operated here by favoring the compensation of sexually dimorphic mechanisms that ultimately guarantees an adequate neuritic development of hypothalamic neurons in both males and females. However, as discussed above, the intriguing assumption that the temporal lag in the development of hypothalamic neurons between males and females could lead to the organization of sex-specific synaptic connectivity patterns *in vivo* remains to be elucidated. A thorough understanding of the molecular and cellular mechanisms that govern early sexual differentiation of the brain will lead to improve our understanding of sex-specific phenotypes in health and disease, allowing us to efficiently address pathologies of embryonic origin that show a clear sex divergence in their manifestations.

**Fig. 9.**
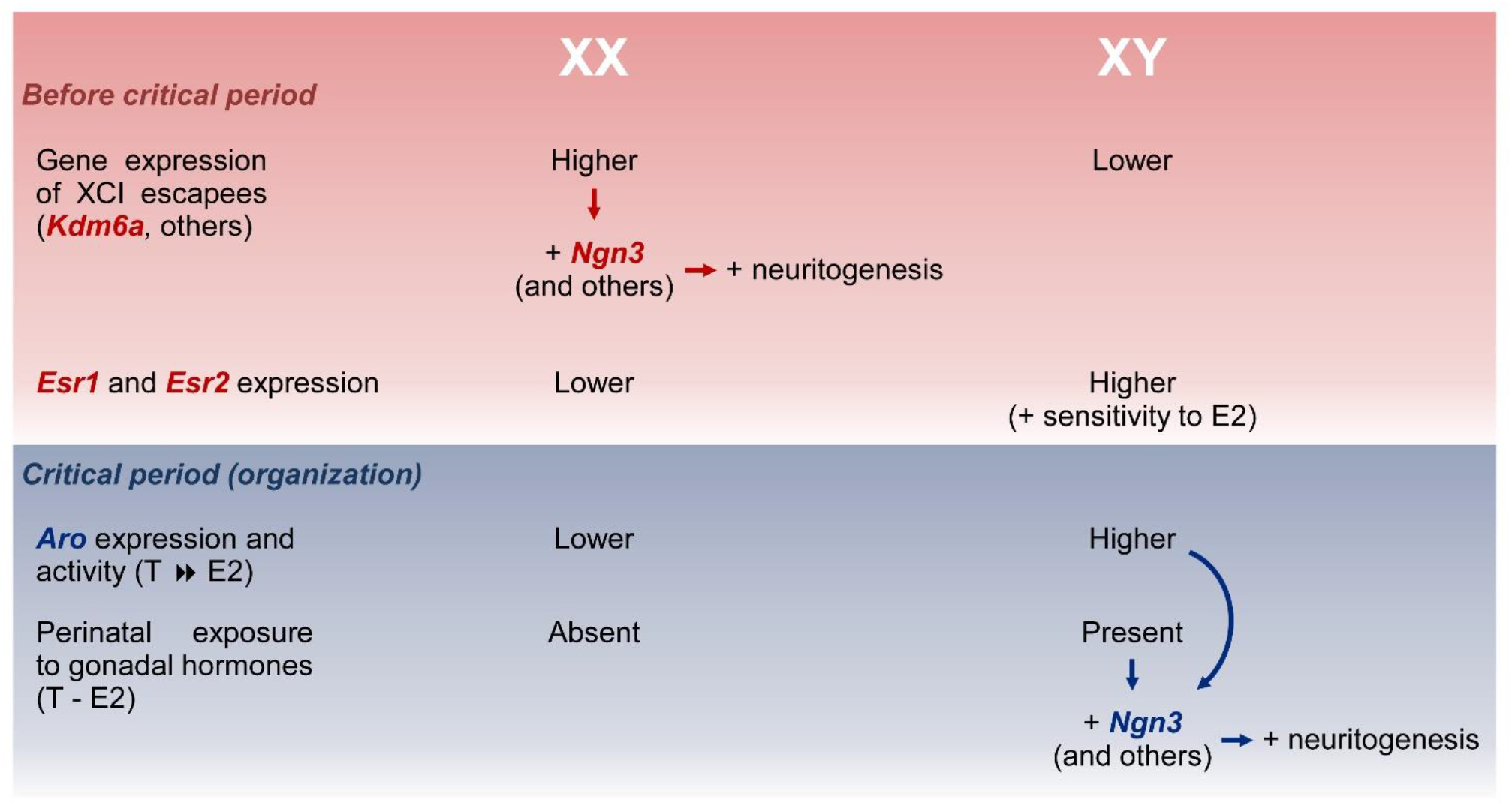
Summary of the hypothesis proposed to explain the sexually dimorphic factors and mechanisms that act in the sexual differentiation of hypothalamic neurons. Before critical period of hormonal organization of the brain, XX neurons show higher expression of X-linked genes that escape X chromosome inactivation (XCI) than XY neurons, such as *Kdm6a*, *Ddx3x*, and *Eif2s3x*. Higher *Kdm6a* levels in XX neurons upregulate neuritogenic genes such as *Ngn3*, *Neurod1*, *Neurod2*, and *Cdk5r1* which, in turn, promotes neuritogenesis. During the critical period, XY neurons are exposed to the effects of estradiol (E2) converted intraneuronally from testosterone (T) by aromatase (Aro), presenting, in addition, a higher expression and activity of this enzyme and a higher sensitivity to E2 than XX neurons by expressing more estrogen receptors *Esr1* and *Esr2*. E2 organizational actions ensure the increase in *Ngn3* expression necessary for the promotion of neuritogenesis in XY neurons.

## Funding

This study was supported by grants from Argentina: Consejo Nacional de Investigaciones Científicas y Técnicas (CONICET, PUE 2016 No. 22920160100135CO), Agencia Nacional de Promoción Científica y Tecnológica (ANPCyT, PICT 2015 No. 1333), and Secretaría de Ciencia y Tecnología de la Universidad Nacional de Córdoba (SECyT-UNC, 2018-2021) to MJC, from Spain: Consejo Superior de Investigaciones Científicas (CSIC, BFU2017-82754-R) to MAA and LMGS and the Enhancing Mobility between Latin America, Caribbean and the European Union in Health & Environment (EMHE)-CSIC Program (MHE-200057) to LMGS and MJC, and from international organizations: International Brain Research Organization (IBRO) Return Home Fellowship and International Society for Neurochemistry (ISN) and Committee for Aid and Education in Neurochemistry (CAEN) Grant to CDC. We would like to thank CSIC and the EMHE Program for financially supporting three research stays of LECZ at the Instituto Cajal, Madrid, Spain.

## Conflicts of interest

The authors declare that they have no conflict of interest.

## Availability of data and material

The datasets generated and/or analyzed during the current study are available from the corresponding author on reasonable request.

## Authors’ contributions

LECZ and MJC conceived and designed the research. LECZ, CDC, and CS performed all experiments and analyzed data. LECZ, CDC, CS, MAA, LMGS, and MJC interpreted results of experiments. LECZ elaborated figures and wrote the first draft of the manuscript. MJC edited and revised the manuscript with critical input from all authors. All authors read and approved the final manuscript. MJC drafted manuscript.

## Ethics approval

All experimental procedures with animals were approved and controlled by the Institutional Animal Care and Use Committee (CICUAL) of the Instituto M. y M. Ferreyra, following national and international regulations, and were in accordance with the Consejería del Medio Ambiente y Territorio (Comunidad de Madrid, Ref. PROEX 200/14), the European Commission (86/609/CEE and 2010/63/UE), and the Spanish Government Directive (R.D.1201/2005) guidelines.

## Notes

### Competing Interest Statement

The authors have declared no competing interest.

